# Linker histone variant H1t is closely associated with repressed repeat-element chromatin domains in pachytene spermatocytes

**DOI:** 10.1101/872408

**Authors:** Iyer Aditya Mahadevan, Sanjeev Kumar, Manchanahalli R. Satyanarayana Rao

## Abstract

**Background:** H1t is the major linker histone variant in pachytene spermatocytes, where it constitutes 50-60% of total H1. This linker histone variant was previously reported to localize in the nucleolar rDNA element in mouse spermatocytes. Our main aim was to determine the extra-nucleolar localization of this linker histone variant in pachytene spermatocytes.

**Results:** We generated H1t-specific antibodies in rabbits and validated its specificity by multiple assays like ELISA, western blot, etc. Genome-wide occupancy studies, as determined by ChIP-sequencing in P20 mouse testicular cells revealed that H1t did not closely associate with active gene promoters and open chromatin regions. Annotation of H1t bound genomic regions revealed that H1t is depleted from DSB hotspots and TSS, but are predominantly associated with retrotransposable repeat elements like LINE and LTR in pachytene spermatocytes. These chromatin domains are repressed based on co-association of H1t observed with methylated CpGs and repressive histone marks like H3K9me3 and H4K20me3 *in vivo*. Mass spectrometric analysis of proteins associated with H1t-containing oligonucleosomes identified piRNA-PIWI pathway proteins, repeat-repression associated proteins and heterochromatin proteins confirming the association with repressed repeat-element genomic regions. We validated the interaction of key proteins with H1t-containing oligonucleosomes by use of ChIP-western blot assays. On the other hand, we observe majority of H1t peaks to be associated with the intergenic spacer of the rDNA element, also in association with SINE elements of the rDNA element. Thus, we have identified the genomic and chromatin features of both nucleolar and extranucleolar localization patterns of linker histone H1t in the context of pachytene spermatocytes.

**Conclusions:** H1t-containing repeat-element LINE and LTR chromatin domains are associated with repressive marks like methylated CpGs, histone modifications H3K9me3 and H4K20me3, and heterochromatin proteins like HP1β, Trim28, PIWIL1 etc. Apart from localisation of H1t at the rDNA element, we demonstrate the extranucleolar association of this linker histone variant at repeat-associated chromatin domains in pachytene spermatocytes. We hypothesize that H1t might induce local chromatin relaxation to recruit heterochromatin and repeat repression-associated protein factors necessary for TE (transposable element) repression, the final biological effect being formation of closed chromatin repressed structures.

## Background

The chromatosome is a structural unit of chromatin, consisting of about 166 bp DNA wrapped around the histone octamer with histone H1 (1, 2). Despite their significant roles in various chromatin-templated biological events, linker histones are not studied in great detail as core histones. H1s possess a unique structure- they contain the N-terminal domain, conserved trypsin-resistant globular domain, and C-terminal domain (3, 4). The N and C-terminal domains of H1 are divergent and largely unstructured in solution (5–7). Its globular domain is the nucleosome binding domain that protects a 20-bp of nucleosomal DNA, just like the full-length H1. On the other hand, the C-terminal domain of H1 is the primary determinant of DNA binding in cells (8). Along with core histones, the linker histone (H1) is one of the five major histone families associated with the eukaryotic chromatin. In mice and humans, eleven H1 variants have been identified to date that includes seven somatic subtypes (H1.0, H1.1-H1.5, and H1x), three testis-specific variants (H1t, HILS1, and H1T2) and one oocyte-specific variant (H1oo).

Mammalian spermatogenesis is an excellent model system to study the biological roles of linker histone variants, as various germ cell variants like H1t, HILS1 and H1T2 are expressed in a stage-specific manner. The testicular linker histone variant H1t is expressed from preleptotene spermatocytes till early round spermatids (in mouse) or till late-round spermatids (in humans) (9–12). Even though H1t mRNA is detected in spermatogonia, the protein is absent. H1t accounts for about 50-60% of total H1 in these cell types (13–15). Surprisingly, the loss of H1t has been shown to cause no detectable defects during spermatogenesis in mice. There are conflicting reports on the phenotypes observed in H1t null mice. For example, H1 subtypes are deposited and been reported to compensate for the loss of H1t (16). In other reports, H1t deficient chromatin is shown to be H1 free (17, 18). Biophysically, H1t is a poor condenser of chromatin in comparison to somatic H1 subtype H1.d (H1.3) as demonstrated by *in vitro* CD (Circular dichroism) spectroscopic studies (19, 20). This property was attributed to a lack of DNA binding motifs like SPKK in the C-terminal domain of H1t (21–24). SPKK and other motifs are responsible for DNA condensation property in histone H1d. Also, in comparison with somatic H1s, a K52Q substitution in the globular domain causes reduction in DNA-binding affinity of H1t (25).

Various defense mechanisms have evolved in germ cells of different species to prevent expression of the retrotransposable elements, thus limiting their mutagenic potential. DNA methylation at LINE and LTR retrotransposable elements can be dependent on piRNA expression or not, also termed as piRNA-dependent and piRNA-independent respectively (26). DNA methylation at LINE retrotransposons are piRNA-dependent (26). In contrast, silencing of LTR retrotransposons by DNA methylation can be piRNA-dependent or independent, with DNA methylation at most of the LTRs are piRNA-independent. Another set of transposable elements like SINE does not exhibit piRNA-dependent repression (27). This suggests that retrotransposon inactivation in germ cells is more complex and their mechanism by the action of important players is currently being investigated in great detail. Retrotranscripts originating from TE (transposable element) sequences in the nucleus are cleaved into sense and antisense piRNAs. The sense RNAs associate with GasZ-containing MILI-TDRD1 complexes within pi-bodies (28). On the other hand, the antisense RNAs associate with MIWI2-TDRD9 complexes and then localize to MAEL-containing piP bodies (29, 30). Degradation of retrotranscripts occurs by the exchange of sense and antisense transcripts in the cytoplasmic compartments. MIWI2-TDRD9 complexes can also induce feedback signaling into the nucleus to facilitate recruitment of Dnmt3L/3A machinery to induce *de novo* DNA methylation at the target TE loci. Some of the essential proteins like MIWI, MAEL, GasZ related to TE repression are also expressed during later stages of spermatogenesis. Mutations in these genes result in pachytene arrest and male infertility.

Our present main aim was to determine the genome-wide occupancy of linker histone variant H1t in pachytene spermatocytes, which would give an idea about its association with specific chromatin domains and consequent biological functions. H1t is not exclusively expressed in testis but also expressed in various cancer cells and mouse embryonic stem cells (31). Recently, H1t-ChIP-sequencing was carried out in human cancer cell lines and mouse ESCs. H1t was found to be majorly associated with the rDNA element of the nucleolus in these cells (31). Also, in the same study, various extra-nucleolar foci of H1t were observed in spermatocytes, which provided major inspiration for the present study to characterize the extranucleolar localization of the linker histone variant H1t in mammalian spermatocytes. It is also important to characterize the binding sites of H1t in the pachytene genome, as H1t is the dominant H1 in these cells, constituting about 50-60% of total H1 content (13–15). This information along with proteins associated with these chromatin domains would give clues towards biological significance of histone H1t in male germ cells due to which they have majorly replaced somatic H1s in the germ cells. We demonstrate that H1t is associated with LTR and LINE repeat element chromatin domains in pachytene spermatocytes. H1t is also closely associated with histone marks H3K9me3 and H4K20me3 and PIWI-piRNA pathway-related proteins suggesting that these chromatin domains represent repressed chromatin domains *in vivo*.

## Results

### Validation of specificity of H1t antibodies

As mentioned earlier, our main aim was to determine the genome-wide localization of linker histone variant in pachytene spermatocytes, for which we required to generate highly specific H1t antibodies. The C-terminal region of H1t is highly divergent compared to other linker histones. The comparison of protein sequences of H1t with somatic linker histone variant H1.2 is given in Fig 1A. As explained earlier, H1t lacks DNA binding motifs that are present in the somatic linker histones (Fig 1A, blue lines). To address the biological functions, we first generated H1t-specific antibodies in rabbits. Since C-terminal domain is highly divergent between H1t and somatic H1s, we cloned and purified the C-terminal protein fragment of H1t (Additional Figure 1A), then used as an antigen to generate polyclonal antibodies in rabbits. We determined the specificity of the antibodies by ELISA and western blotting assays. By ELISA assays, we found that the sera, as well as purified antibody, reacted to the recombinant H1t C-terminal protein fragment (Additional Figure 1B and Additional Figure 1C respectively). Western blotting also showed specific reactivity with the protein corresponding to H1t in the testicular nuclear lysates (Fig 1B, α-H1t lane). Also, we observe a different band when H1.2 antibodies were used (Fig 1B, α-H1.2 lane), suggesting that the molecular weights of the bands were specific to these variants and did not crossreact with other variants. We also confirmed the reactivity of H1t antibodies against mouse testicular histones by western blotting and obtained the specific band corresponding to H1t (Fig 1C). This was additionally confirmed by mass spectrometry, as H1t was the only variant to be associated with H1t-containing oligonucleosomes (See later). We further determined the staining pattern of H1t across the various stages of meiotic prophase I. We observed uniform distribution of H1t protein across the leptotene, zygotene and pachytene cells (Additional Figure 1D), consistent with being the dominant H1 in spermatocytes. The combination of western blotting, ELISA and mass spectrometry assays thus establish the specificity of the in-house generated H1t antibodies.

**Figure 1.**
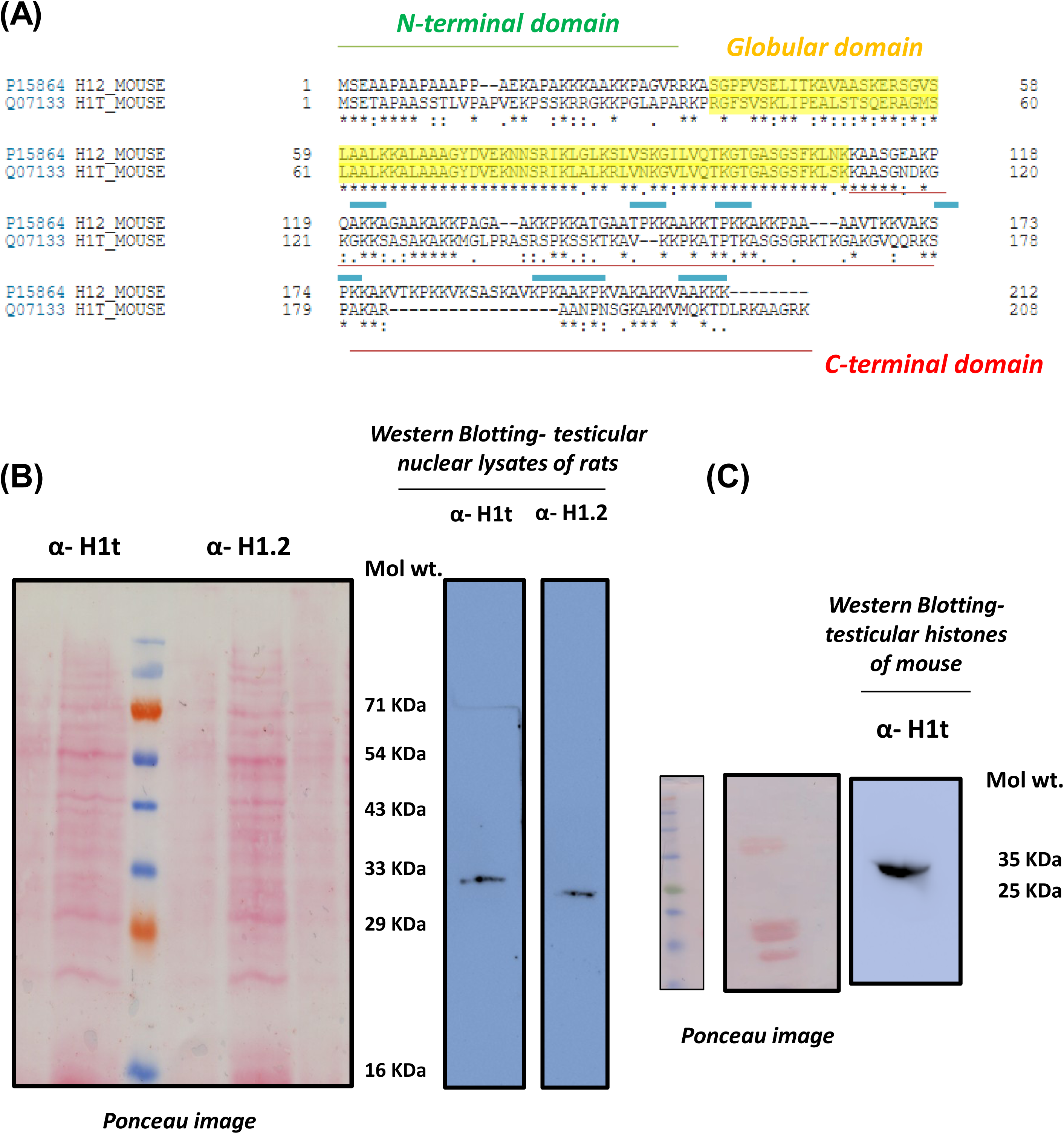
Generation and validation of specificity of H1t-specific antibodies. **A.** The tripartite structure of linker histones- the linker histones possess three major domains: N-terminal domain, globular domain, and the C-terminal domain. The DNA binding motifs like SPKK that are present in the somatic linker histone H1.2 have been underlined using blue lines. Since the C-terminal of H1t protein is highly divergent in comparison with other variants, we used 112-207 amino acid residues as protein fragment for the generation of H1t-specific antibodies in rabbits. **B.** Western blotting of H1t and H1.2 antibodies against nuclear lysates prepared from P25 rat testicular cells. The anti-H1t and anti-H1.2 antibodies showed reactivity to the specific linker histones, as seen in the blot images, where bands corresponding to different molecular weights were obtained. Ponceau stained blots are given for reference. **C.** Western blotting of H1t antibodies against P20 mouse testicular histones. Ponceau stained blots are given for reference.

### Genome-wide occupancy of linker histone variant H1t in pachytene spermatocytes

Since H1t is a linker histone and a component of chromatin, we carried out ChIP-sequencing with the crosslinked chromatin to determine the occupancy sites of H1t in the pachytene chromatin of mouse. The experimental workflow of the ChIP protocol is given in Figure 2A. The profile of DNA fragments obtained after various cycles of sonication is given in Additional Figure 2A. We chose mouse testis as the model system for the study, since various datasets related to biological processes like meiotic recombination, transcription have been well characterized in the mouse species.

**Figure 2.**
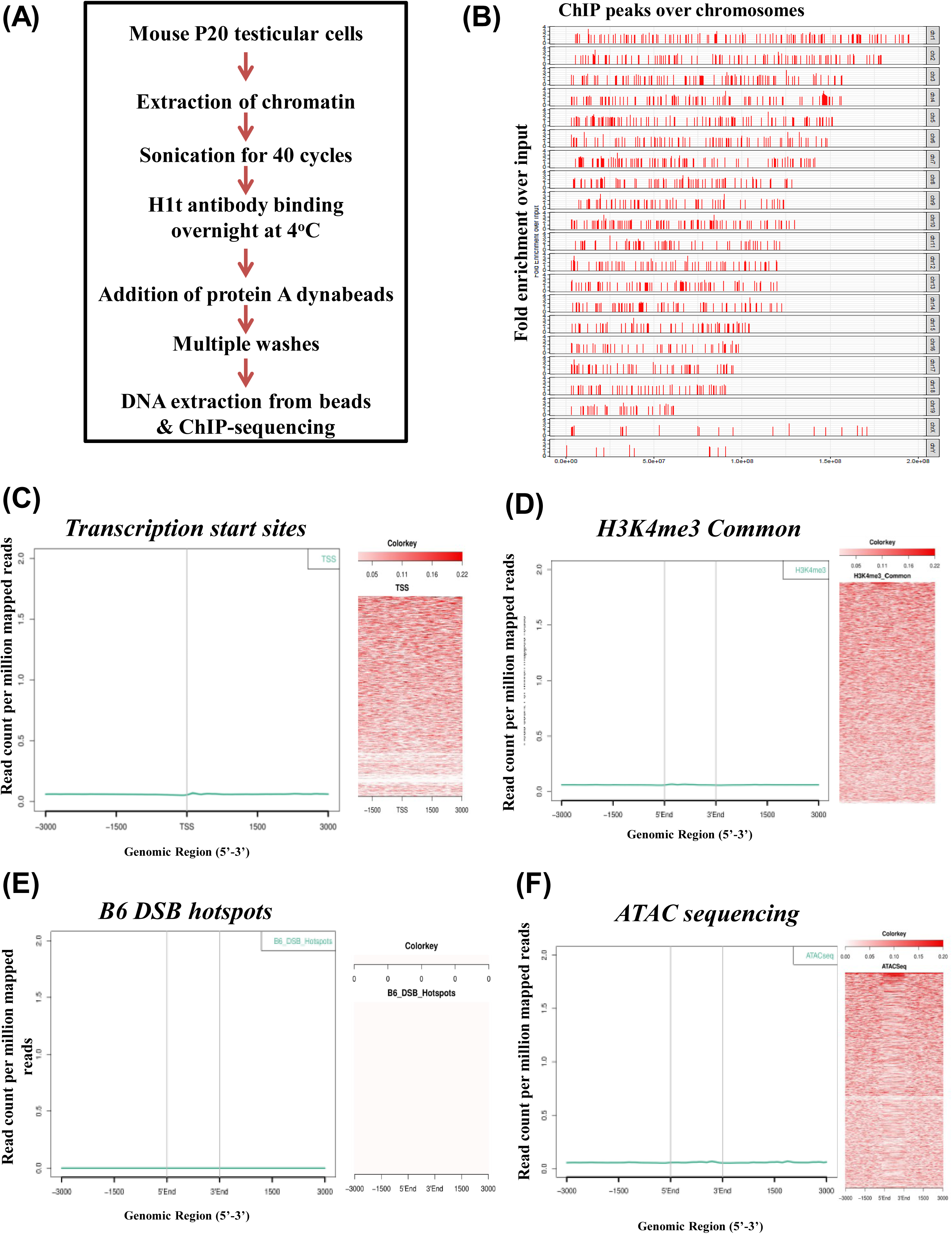
Localization of linker histone variant H1t at TSS, active gene promoters, recombination hotspots and open chromatin regions (ATAC seq positive regions) **A.** Workflow of the experimental protocol used for the H1t ChIP-sequencing technique. **B.** Chromosome-wise distribution of H1t peaks in the pachytene genome. The y-axis represent the fold enrichment observed for H1t IP over background input, and the x-axis are the various H1t-occupied locations along the length of each chromosome. The normalisation of high fold enriched peaks has been done to 3.5 fold change and the lower threshold has been maintained for fold change=2. Analysis of overlap between H1t and **C.** TSS of the mouse (GENCODE), **D**. H3K4me3 common representing the TSS-associated H3K4me3 marks (GSE93955), **E.** DSB hotspots (GSE93955), **F**. Open chromatin regions of pachytene spermatocytes (32). H1t were observed to depleted at active gene promoters, TSS regions, and open chromatin genomic regions. Information in (**C-F**), the overlap has been determined using aggregation plots (left panels) and heat maps (right panels).

By ChIP-sequencing and the related data analysis, we obtained statistically significant (p value <= 0.05) 48681 peaks of H1t occupancy (Refer methods section for details). The chromosome-wise distribution of these H1t peaks is shown in Figure 2B. To reiterate, linker histones are known to be generally depleted from active TSS, open chromatin structures *in vivo*, except some variants like H1x, H1.1 (Izzo et al., 2013). We wanted to determine whether the variant H1t is associated with genomic regions related to transcription, meiotic recombination, etc. We performed overlap analysis of H1t genomic peaks with other ChIP-sequencing datasets and the results are represented as aggregation plots and heat maps. Both these methods show the spatial distribution of reads within target genomic regions. Aggregation plots as given in Fig 2C and 2D (left panels), shows that H1t is not significantly associated with transcription start sites as well as H3K4me3-marked active gene promoters. The heat maps corroborate with this data as no significant overlap was observed for H1t reads at TSS and active gene promoters (Fig 2C, 2D, right panels). When analyzed with DSB hotspots, we observe H1t also not to be closely associated with recombination hotspot genomic sequences (Fig 2E). Furthermore, when performing the overlap analysis of H1t ChIP-seq dataset with ATAC-sequencing dataset available for pachytene spermatocytes (32), we observed that H1t is not majorly associated with open chromatin regions (ATAC-seq positive genomic regions) in the pachytene genome (Fig 2F). These observations suggested to us that H1t might be associated with chromatin regions whose functional consequence would be to ultimately form condensed chromatin structures *in vivo*.

Since H1t is not associated with TSS, active gene promoters, DSB hotspots, and ATAC-seq positive regions, the primary question remained to what genomic regions are H1t localized at. Upon initial annotation of the H1t-occupied genomic regions, we observed that H1t is majorly associated with intergenic and intronic regions (Fig 3A). On further annotation of the H1t-bound genomic regions, we observed that the majority of the H1t were closely associated with LINE and LTR classes of repetitive elements, and SINE to a lesser extent (Fig 3B). We therefore conclude that H1t is localized to retrotransposable elements LINE, LTR and SINE *in vivo*. It is now well established that these retrotransposable elements are repressed in the germ cells by the action of RNA interference machinery and small RNAs. Despite millions of years of divergence, the machinery involving RNAi machinery and piRNAs in preventing transposable element (TE) expression have been conserved in fungi, plants, and animals. piRNAs are essential for de novo DNA methylation at these TE loci crucial for silencing of LINE and LTR elements in embryonic male germ cells. DNA methylation is carried out by the methyltransferase enzyme Dnmt3L in early germ cells, the loss of which results in male infertility due to meiotic failure (33). Therefore, DNA methylation of repeat elements is a major mechanism for preventing TE expression and is critical for success of productive spermatogenesis. Since the piRNA pathway machinery repress TE by DNA methylation, our next question was to determine whether H1t-associated genomic regions are also associated with methylated genomic sequences. We carried out the overlap analysis of our H1t ChIP-sequencing data with the already published bisulfite sequencing dataset available for P20 mouse testicular cells (34). As can be seen in Fig 3C and Fig 3D, we observed more than 90% (44720 peaks) of H1t peaks were associated with methylated CpGs at these repetitive elements. These observations suggested that H1t, apart from its major association with classes of repetitive elements LINE and LTR elements that are methylated in pachytene spermatocytes.

**Figure 3.**
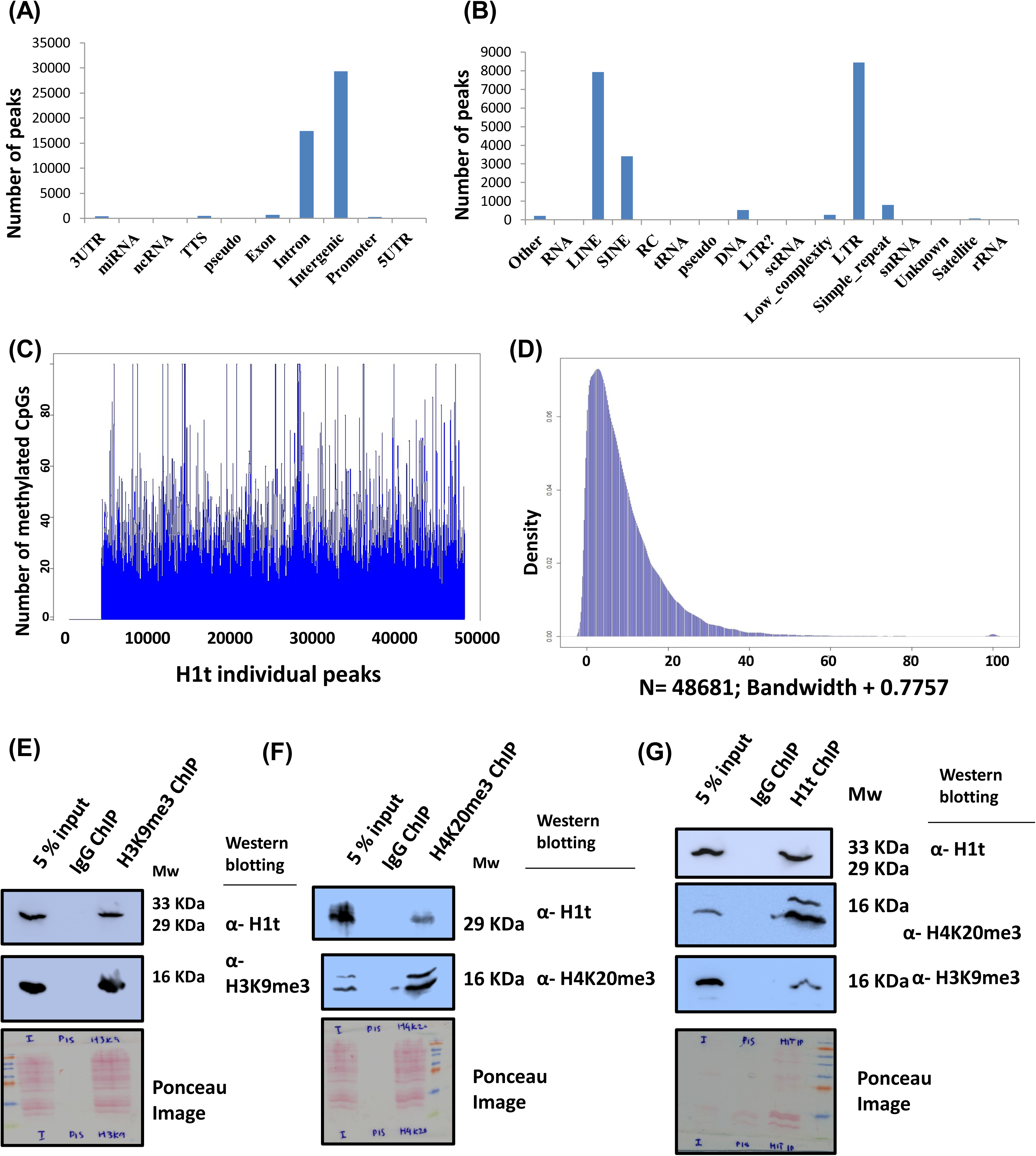
Localization of linker histone H1t at CpG-methylated repeat element chromatin domains. **A.** Annotation of H1t-bound genomic regions using HOMER **B.** Annotation of H1t peaks at repeat elements showing its predominant association with LINE and LTR subclasses of retrotransposable elements. **C.** Profile of methylated cytosines (SRS557654) across all the H1t peaks. More than 90% of the H1t peaks overlap with methylated CpGs. The y-axis is the count of methylated CpG positions at each peak. Some peaks with more than 100 methylated cytosines have been truncated to 100 for better visualization of the overall plot. **D.** Density plot of H1t peaks overlapping with methylated CpGs (SRS557654). The y-axis represents the density function and the x-axis represents the bandwidth parameter. N represents the number of observations. **Co-immunoprecipitation assays showing the coexistence of H1t-containing oligonucleosomes with histone marks H3K9me3 and H4K20me3.** **E.** H3K9me3-ChIP showing the co-association with linker histone variant H1t in testicular chromatin. **F.** Linker histone variant H1t is associated with H4K20me3-ChIP elute fraction **G.** Reciprocal immunoprecipitation assays showing that H1t-positive chromatin fragments associated with histone marks H3K9me3 and H4K20me3 *in vivo*. Information in (**E-G**) The first lane is the input fraction; the second lane is the IP using the non-specific IgG isotype control, the third lane is the IP with the mentioned antibodies (anti-anti-H3K9me3/anti-H4K20me3/anti-H1t). The antibodies labeled alongside the blot refers to the antibodies used for western blotting. Ponceau stained blots are given for reference.

In addition to DNA methylation, repressive histone modifications like H3K9me3 (35) and H4K20me3 (36) facilitate silencing of the retrotransposable elements. H3K9me3 histone mark is present on LINE and LTR repeat elements in germ cells and is dependent on the function of piRNA pathway (35). H3K9 methylation is an important epigenetic mark in transcriptional silencing and heterochromatin formation. H3K9me3 formation is mediated by histone methyltransferases (HMTs) Suv39h1 and Suv39h2 in germ cells (37–39). We went ahead to determine whether H1t-containing oligonucleosomes are associated with repressive histone marks like H3K9me3 and H4K20me3 *in vivo*. By employing oligonucleosome IP assays, we found that H3K9me3 and H4K20me3 did pull down H1t protein (Fig 3E, 3F, respectively). Also, by reciprocal IP assays, we observed H1t-associated oligonucleosomes to be positive for H3K9me3 and H4K20me3 histone marks *in vivo* (Fig 3G). These biochemical assays demonstrate the association of repressive histone marks H3K9me3 and H4K20me3 with H1t-bound chromatin fragments in pachytene spermatocytes. Thus, we provide strong evidence to the fact that H1t-containing genomic regions are associated with DNA methylation and repressive histone modifications H3K9me3 and H4K20me3 in pachytene spermatocytes.

### Localization of linker histone variant H1t in the rDNA element of the pachytene spermatocyte

Linker histone variant H1t was previously demonstrated to be localized to the rDNA element of the nucleoli in spermatocytes by immunofluorescence assays (31). We confirmed the association of H1t in the rDNA element of pachytene spermatocytes, wherein we observe 13008 peaks to be localized at the known mouse rDNA element. Interestingly, the majority of the H1t peaks are localized in the intergenic spacer of the rDNA element (Fig 4A).

**Figure 4.**
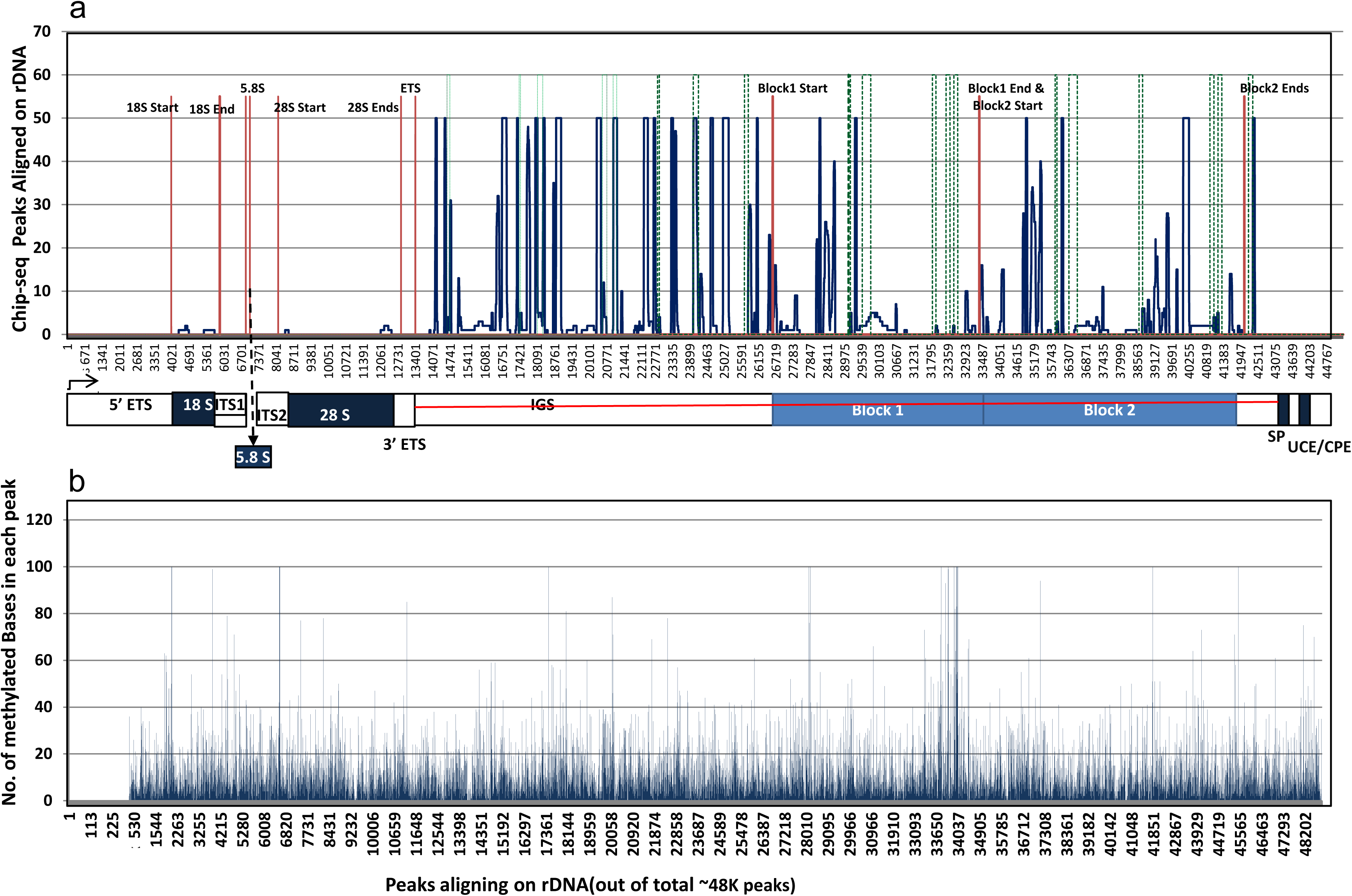
Localization of linker histone H1t in the mouse rDNA element. **A.** Peak distribution of H1t across various regions of the rDNA element (40). Distribution of H1t peaks with respect to SINE elements (green lines) localized in the rDNA element. **B.** Distribution of methylated CpGs across all the H1t peaks in the rDNA element. More than 99% of the H1t peaks overlap with methylated CpGs in the rDNA element. The y-axis represents the number of methylated CpGs, and the x-axis represents the individual H1t peaks that are localized in the rDNA element. Information in **(A-B)** The red lines demarcate various regions of the rDNA element (40), blue lines are the H1t peaks, and the various regions of the rDNA element have been labeled below the peak distribution maps.

The rDNA element harbors many repetitive elements, the dominant element being SINE (40). SINE elements constitute about 20% of the total rDNA element in the mouse. We observe that H1t peaks are located at or close to the vicinity of the predominant SINE elements (marked in green) in the rDNA element of the pachytene spermatocyte (Fig 4A). We also wondered whether the H1t peaks localized at the repetitive elements of the rDNA element are associated with DNA methylation. True to our intuition, we observed more than 95% (12550 peaks) of H1t peaks are indeed associated with methylated CpGs at the rDNA element (Fig 4B). In addition to the extra-nucleolar localization of H1t at the retrotransposon classes LINE and LTR, we also observe significant overlap of H1t at the repressed repetitive elements of the rDNA element, characterized by occupancy of methylated CpGs. Thus, in both the nucleolar and extranucleolar mouse genome, linker histone variant H1t occupies the methylated CpG repeat-associated chromatin domains.

### Mass spectrometric identification of proteins associated with H1t-bound chromatin fragments

H1t being the major linker histone component of pachytene chromatin, we wanted to identify the proteins that co-associate with H1t-containing chromatin fragments. For this purpose, after carrying out the H1t ChIP, we performed mass spectrometric analysis of the eluted proteins, to identify the H1t-associated proteins in the chromatin context. The proteins were identified based on the enrichment of proteins observed in the H1t ChIP fraction compared to non-specific rabbit IgG ChIP fraction (Fig 5A). Interestingly, we observed H1t be the only linker histone variant associated with the H1t ChIP fraction. This provides additional strength for our studies regarding the specificity of the H1t antibody, highlighting the robustness of our observations.

**Figure 5.**
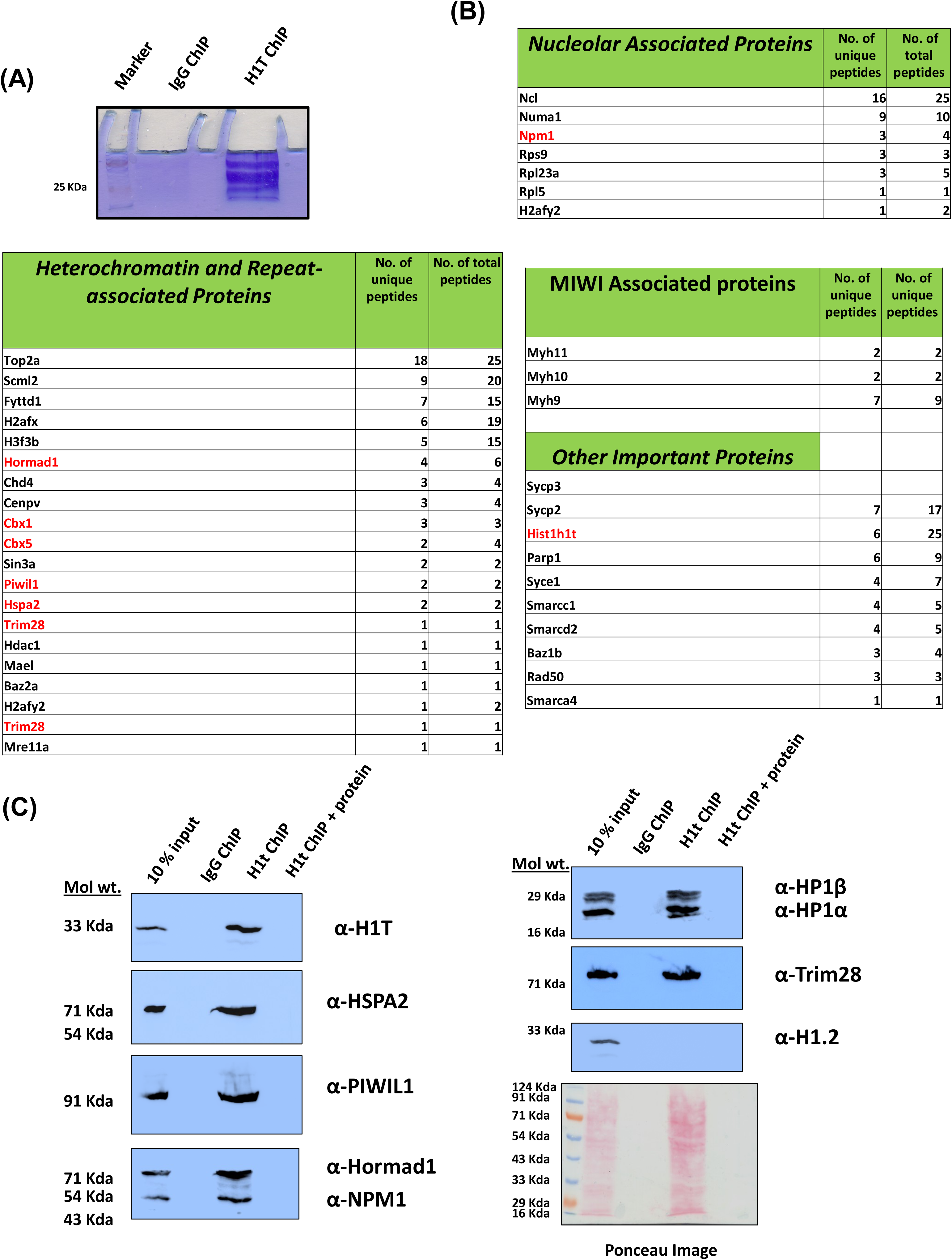
List of key proteins associated with H1t-positive chromatin fragments in pachytene spermatocytes as determined by mass spectrometry. **A.** Coomassie-stained gel showing the success of the H1t-ChIP technique. The first lane represent the ladder, the second lane is the ChIP carried out using the rabbit IgG antibodies and the third lane is the ChIP carried out using H1t antibodies. **B.** The important proteins that are associated with H1t-containing oligonucleosomes can be divided into four major classes: Nucleolar function, heterochromatin and repeat-associated proteins, MIWI associated proteins and Other important proteins. The proteins indicated in red color have been selected for further validation by co-IP assays. **C.** Validation of H1t-associated proteins by ChIP-western blotting technique- To validate further the association of key proteins with H1t-associated chromatin, we carried out western blot analysis of the H1t-positive oligonucleosomes. We observe proteins related to the nucleolus (NPM1), heterochromatin related (Hormad1, Trim28, HP1β, HP1γ), PIWI pathway-related (PIWIL1, Hspa2) are associated with H1t-containing ChIP fraction. The first lane in all the blots represents the 10% input fraction, second lane ChIP with the non-specific isotype control, and the third lane ChIP with the H1t antibody, and the fourth lane is the ChIP carried out along with addition of C-terminal H1t recombinant protein (10µg). The antibodies labelled in alpha alongside the blot represent the antibodies used for western blotting.

Further, we observed that the H1t-associated proteins could be identified into three classes-nucleolar-associated, repeat element and heterochromatin-associated and other important proteins (Fig 5B). We expected association of nucleolus related proteins as H1t is shown to be localized at the rDNA element of mouse spermatocytes (Fig 5B, nucleolar proteins) (31). Importantly, we observed various PIWI-piRNA pathway proteins such as Piwil1(MIWI) and its associated proteins (Myh9, Myh10, Myh11), HSPA2, MAEL to be associated with H1t-oligonucleosomes (Fig 5B, repeat-associated and heterochromatin proteins, MIWI associated proteins). Piwil1 protein is a bonafide slicer (small-RNA-directed endonuclease) in postnatal germ cells wherein the loss of Piwil1 has been shown to cause male fertility due to the upregulation of LINE1 retrotransposon transcripts (41). Also, Tdkrh1 is known to interact with PIWI proteins and is an essential factor implicated in piRNA biogenesis (42). These results provide additional evidence to the association of H1t with proteins related to TE silencing in pachytene spermatocytes. The entire list of proteins obtained after mass spectrometry is given in Additional File 3. Also, the list obtained on comparison of H1t-associated proteins with the comprehensive published list of heterochromatin proteins (43) is given in Additional File 4. Further, we have validated the association of H1t with some of the key proteins by carrying out ChIP assays followed by western blotting (using the antibodies for detection of proteins highlighted in red), and observed that H1t is associated with proteins related to nucleolar function (Npm1), repeat-repression and heterochromatinization (HP1β, HP1α, Piwil1, Hspa2, Hormad1, Trim28) *in vivo* (Fig 5C, third lanes). H3K9me3-positive nucleosomes are the docking sites for the binding of chromodomain-containing HP1 proteins establishing the chromatin condensed structure. This chromatin template has been proven for transcriptional repression, also leading to spreading and amplication of heterochromatin domain. The fact that H1t-containing nucleosomes associate with H3K9me3, HP1α, HP1β gives a strong indication for the localization of linker histone H1t at repressed repeat-element chromatin domains.

In addition to H3K9me3 and H4K20me3, there could be other histone marks that could be signatures of repeat element nucleosomes. Uhrf1 has been shown to interact with PRMT5 (arginine methyltransferase) to catalyze the formation of H4R3me2s and H3R2me2s. These marks also act together with PIWI proteins to facilitate retrotransposon silencing in the germline (44). We found that H1t-oligonucleosomes associate with Tdkrh, a known interacting protein partner with Uhrf1 and PIWI proteins, further supporting our observations of association of H1t with repressed-repeat element chromatin domains in pachytene spermatocytes. Importantly, preincubation of the H1t-antibodies with C-terminal protein of H1t blocked the reactivity of the antibodies, reconfirming the validity of our observations (Fig 5C, fourth lanes). We further observed that the major somatic linker histone H1.2 not to be associated with H1t-ChIP elute fraction, highlighting that the H1t-antibodies did not pull down H1.2 and its associated proteins, but were specific to the linker histone variant H1t (Fig 5C, α-H1.2, third lane). Mass spectrometric analysis of H1t-associated protein complexes reflect the average picture at all the repeat-element chromatin domains. Repeat-class specific complexes could exist to modulate loci-specific functions in pachytene spermatocytes. In summary, these observations demonstrate the association of linker histone H1t with repressed repeat-chromatin domains in pachytene spermatocytes.

## Discussion

Recently, tremendous progress has been made towards understanding the molecular functions of linker histone H1 and its subtypes in developmental processes beyond their architectural roles in chromatin. The lack of studies on H1 function can be largely attributed to technical difficulties during the use of mass spectrometry approaches yielding reduced protein coverage and also the generation of subtype-specific antibodies. Raising variant-specific antibodies is a challenge. In this background, we have successfully generated H1t-specific antibodies in rabbit and validated its specificity by multiple assays.

### Linker histone variant H1t and TE repression

An important observation made in the present study is the predominant localization of linker histone variant H1t at repeat elements belonging to LINE and LTR. Even though repeat element RNAs orchestrate gene expression patterns during early embryonic development (45, 46), paradoxically in the context of germ cells, the repeat elements need to be repressed to prevent mutagenesis and genome instability. Importantly, the expression of proteins like Dnmt3l, the enzyme that is responsible of DNA methylation of repeat elements, does not occur in spermatocytes, establishing methylation patterns in prespermatogonia become necessary in spermatocytes, because the defects are inherited during the pachytene interval. The loss of Dnmt3l leads to the upregulation of LINE and LTR transcripts in spermatogonia and spermatocytes, suggesting its role in TE repression phenomenon in the premeiotic germ cells. These epigenetic processes involving TE repression are important especially during pachytene interval, where scanning and apoptotic checkpoint mechanisms are in place. Importantly, defects in DNA methylation at TE elements results in shifting of recombination hotspots to non-canonical genomic sites, ultimately resulting in male infertility (47). Based on the observation of the H1t-occupied repeat elements coinciding with occupancy of methylated CpGs, it is tempting to speculate that these repeat elements represent repressed chromatin domains *in vivo*. H1 has also been shown to contribute to the establishment of DNA methylation patterns in mESCs (48); this provides additional support for the importance of H1t and its association with DNA methylation-associated genomic regions in pachytene spermatocytes. In *Arabidopsis thaliana*, linker histone H1 and DNA methylation jointly regulate gene expression patterns by repressing TE *in vivo* (49). Thus, it is very likely that even in the context of mammalian spermatocytes, H1t and methylated CpGs might recruit specialized effector proteins to jointly repress TE chromatin domains (6A, 6B).

An interesting question that arises here is what determines the localization of H1t to these specific genomic loci. We observed a combination of twelve motifs to be significant in the total H1t peaks as identified through motif analysis using MEME (Additional Figure 2B). Whether the motif information of the target DNA sequence is sufficient for H1t binding in the pachytene spermatocyte genome remains to be further examined. Since we observed that the occupancy of the repeat elements is characterisitic of linker histone H1t in pachytene spermatocytes, it could be possible that the unique C-terminus of H1t carry information to localize at repetitive elements. It is also possible that adaptor proteins might recruit histone H1t to these specific genomic loci. In this context, we would like to mention that PIWI proteins recruit H1 to TE loci to repress these chromatin domains in ovarian somatic cells (OSC) of Drosophila (50). Therefore, proteins belonging to the PIWI pathway might regulate H1t localization at repeat elements in pachytene spermatocytes. This presumption is additionally supported by the fact that various proteins belonging to PIWI pathway like PIWIL1, Hspa2, Uhrf1, Trim28 are associated with H1t-containing chromatin domains. It is also known that C-terminal domain of H1 is required for recruitment of Piwi and TE silencing in Drosophila (50). We were surprised to see that the Drosophila H1 and H1t protein sequences are similar in the C-terminal regions being devoid of various DNA binding motifs like S/TPKK (Fig 6D).

**Figure 6.**
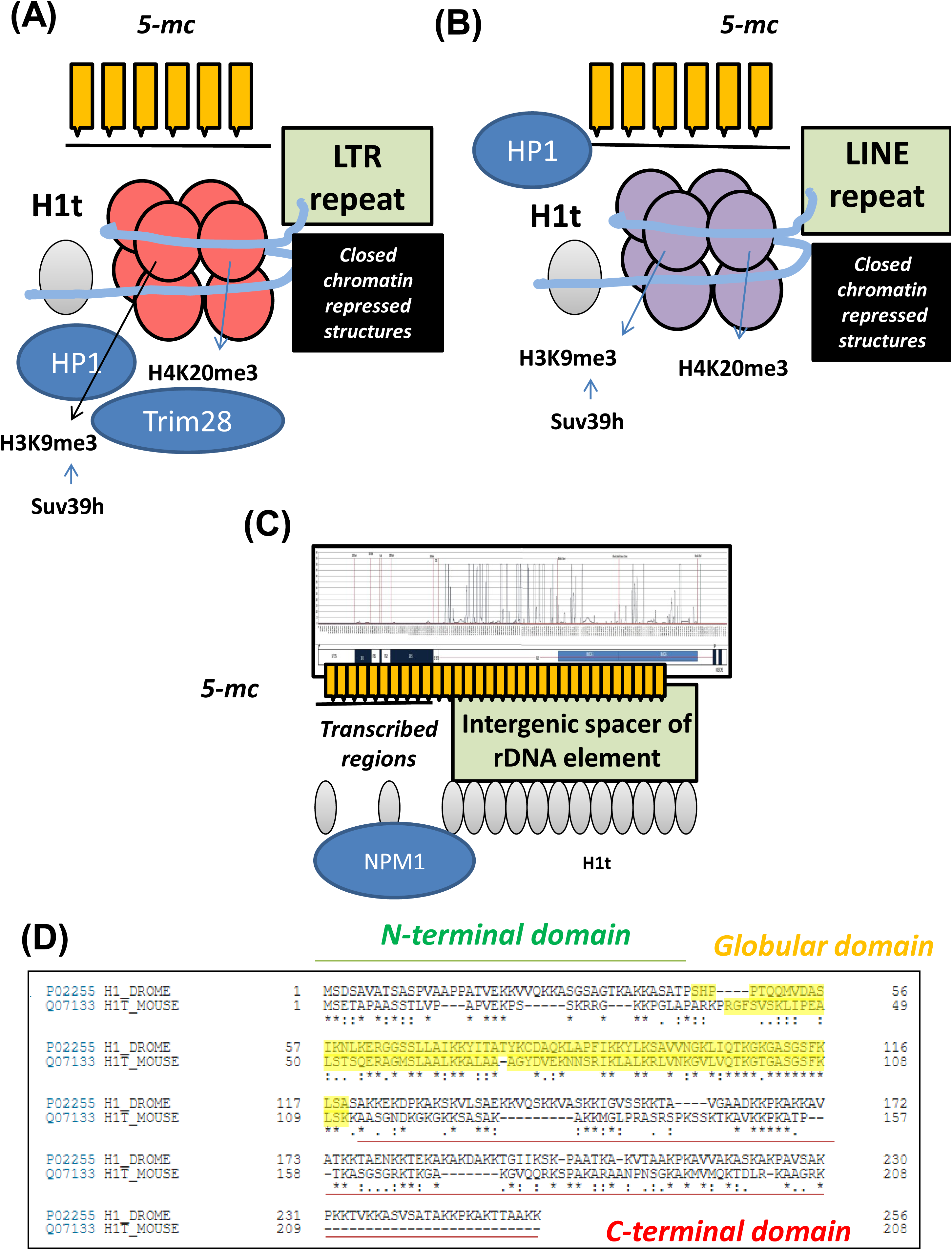
Model of H1t-containing chromatin domains in pachytene spermatocytes. **(A-B)** Extra-nucleolar localization of linker histone variant H1t is related to **A**. LINE and **B**. LTR repeat-element chromatin domains. This forms a chromatin template for binding of PIWI proteins, DNA methylation machinery (forming methylated CpGs), repressive histone modifications like H3K9me3 and H4K20me3 in pachytene spermatocytes. H3K9me3 nucleosomes are binding sites for HP1 proteins to mediate heterochromatinization at these target loci. **C.** Predominant association of linker histone variant with the intergenic spacer of the rDNA element. H1t co-associates with methylated CpGs and nucleolar proteins like Npm1 at the rDNA element in pachytene spermatocytes. **D.** Alignment of protein sequences of mouse H1t with Drosophila H1. Both these proteins lack DNA binding motifs like S/TPKK in its C-terminal domain.

H1t-containing chromatin domains are also associated with repressive histone modifications like H3K9me3 and H4K20me3 in the spermatocyte genome. It will be interesting to decipher whether the association of H1t with methylated CpGs in the repeat elements is dependent on the piRNA-PIWI pathway or not. Despite ongoing efforts to characterize in-detail the mechanisms of TE repression in germ cells, the crosstalk between epigenetic pathways involving repressive histone modifications, DNA methylation, and piRNAs/PIWI proteins have not been delineated in great detail. We suspect H1t along with repressive histone H4K20me3 and H3K9me3 might set the template for TE repression in germ cells (Fig 6A, 6B). Crosstalk between H1t and repressive histone modifications could occur; working with conjunction with piRNA-PIWI pathway and heterochromatin proteins can lead to formation of repressed chromatin structures *in vivo*.

Biochemical and biophysical studies have shown that linker histone H1t, due to lack of S/TPKK motif in its C-terminus, is a poor condenser of chromatin and also generates a relaxed chromatin structure with oligonucleosomal template *in vitro* (19–24). Based on this propensity, we would like to propose that H1t-mediated relaxed chromatin structure at the repeat element chromatin domains would facilitate the initiation of recruitment of repression machinery, resulting in heterochromatin formation at these sites.

### H1t and nucleolus

As explained earlier, H1t was previously shown to be localized to the nucleolus in mouse spermatocytes and human cancer cell lines (31). We could confirm their observations even in spermatocytes. Further evidence for association of linker histone H1t with nucleolar chromatin comes from our observations that we could pick up nucleolar proteins such as Npm1, Numa1, Ncl in the H1t IP chromain by mass spectrometry. An interesting observation made in the present study is the predominant association of H1t peaks with the intergenic spacer of the rDNA element. There have been numerous reports of the role of intergenic spacer in modulating rDNA transcription (51, 52). macroH2A has been shown to repress rDNA transcription in human HeLa and mouse ES cells (53). Since we observed macroH2A be associated with H1t-ChIP fraction as determined by mass spectrometry, this suggests that H1t could function with macroH2A to modulate rDNA transcription in germ cells, providing further support to the possible role of H1t in rDNA transcriptional dynamics. The rDNA element is a host to repetitive elements of various classes, SINE being the predominant class (40).

Both the nucleolar and extra-nucleolar localization of H1t occurs in the repeat regions of the mouse pachytene genome. In mammalian cells, chromatin is organized into structural and functional compartments like TADs (topologically associated domains), LADs (lamin associated domains) etc (54–59). Recently, nucleolus associated domains (NADs) have been discovered in HeLa cervical carcinoma and HT1080 fibrosarcoma cells (60, 61). Various repeat elements like LTR are enriched in NADs and inter NADs (62). It would be interesting to see whether this kind of nucleolar organization exists in pachytene spermatocytes, if yes, then does H1t influence structure of NADs and inter NADs via its association with repeat elements.

Nucleosomal retention occurs in repetitive sequences associated with intergenic and intronic regions in mammalian sperm (63, 64). This epigenetic landscape sets up an important marker for paternally derived nucleosomes in preimplantation embryos. Since H1t has been demonstrated to occur at repeat sequences associated with intergenic and intronic regions, this begs for the question of what linker histone variants or PTMs mark these functionally important genomic regions in the mature sperm. An important point to be noted here is that H1t expression is restricted till early round spermatids in mouse, other variants at repeat elements could replace H1t during spermiogenesis. One candidate variant, HILS1 is also enriched at the LINE1 elements in rat spermatids (65). Also, *in vitro* studies have demonstrated that H1t can be poly ADP-ribosylated (PAR), and PAR modified H1t promotes chromatin compaction (66). Disturbance in PAR metabolism causes retention of H1t, HILS1 and core histones in the mature spermatozoan (67). Recently, various post-translational modifications have been characterized on H1t obtained from spermatocyte and round spermatocyte cell populations by mass spectrometry (68). It would also be interesting to see whether PTMs on H1t mark specific repeat element subclasses. It will be exciting to determine the loci-specific functions of H1t PTMs in repeat repression phenomena. Specific marking of chromatin territories might occur in various stages of germ cell development, wherein different histone variants with their PTMs might contribute to unique functions in shaping the epigenetic landscape important for male fertility and transgenerational inheritance. An intriguing question that arises at this juncture is why histone H1t knockout mice do not show any defect in fertility. It is worth mentioning here that the histone TH2B knockout mice are also fertile, but in these mice, the somatic histones H2B, H3 and H4 acquire compensatory histone PTMs that could subsitute the role of TH2B in spermatogenesis (69). Thus, a more detailed investigation of chromatin modifications in histone H1t-deficient mice is necessary to understand the biological function and relevance of histone H1t in mammalian spermatogenesis.

## Conclusions

Very few studies have shed light on the biological roles of testicular linker histone variants in the context of differentiation of male germ cells. The present study is focussed towards understanding the genomic and chromatin features of H1t-occupied chromatin domains. We observe that linker histone H1t is not closely associated with TSS, active gene promoters, DSB hotspots and open chromatin regions. We have extensively characterized the nucleolar and extranucleolar localization features of linker histone variant H1t and found that H1t to be predominantly associated with repeat element chromatin domains in pachytene spermatocytes. These chromatin domains are positive for methylated CpGs, repressive histone modifications like H3K9me3 and H4K20me3, protein marks like piRNA-PIWI pathway components, heterochromatin proteins etc. We propose that H1t might induce local chromatin relaxation to recruit proteins effectors necessary for repression of these repeat elements.

## Materials and Methods

### Cloning and expression of C-terminal protein fragment of H1t

The coding sequence (CDS) corresponding to 112-207 amino acid residues of H1t protein was cloned using specific primers (Forward primer-ATAGAAGCTTGCAGCTTCAGGCAACGAC and Reverse primer-ATATGCGGCCGCCTTTCTTCCTGCTGCCTTCC) into pET22b(+) vector using HindIII and NotI restriction sites. The expression vector was transformed in Rosetta strain of E.coli, and His-tagged proteins were purified using Ni-NTA purification method.

### Antibody generation

The recombinant protein (C terminal fragment of H1t) was injected into rabbits, and the 14-day cycle of antibody generation was followed. Immunoglobulins were purified by caprylic acid-based purification method. Antigen-affinity based purification with the Sulfolink columns containing immobilized proteins was used to purify the H1t-specific antibodies. The H1t antibody was outsourced from the Abgenex company (Bhubaneshwar, India).

### ELISA

The recombinant proteins were used at 200 ng per well. The pre-bleed and immune sera were used at 1:5000 dilution. Goat anti-rabbit HRP was used as the secondary antibody at 1:5000 dilution. TMB (3, 3’, 5, 5’-Tetramethylbenzidine) was used as the substrate for color development. After three minutes of enzyme-substrate reaction, the plate was read at 450 nm.

### Preparation of testicular nuclear lysates

Nuclear lysates were prepared by the method described previously with modifications (70). Briefly, testes were dissected in cytoplasmic lysis buffer (10mM HEPES pH 7.5, 50mM NaCl, 0.5M sucrose, 0.5% Triton-X-100, 0.1mM EDTA, 1mM DTT, protease inhibitor cocktail), incubated on ice for 15 minutes and centrifuged at 1500g for 7 minutes. The nuclear pellet was resuspended in Buffer B1 (10mM HEPES pH 7.5, 500mM NaCl, 0.1mM EDTA, 1mM DTT, 0.5% NP-40, protease inhibitor cocktail) to obtain nuclear lysates or Buffer B2 (10mM HEPES, 200mM NaCl, 1mM EDTA, 0.5% NP-40, protease inhibitor cocktail) for isolation of chromatin. The nuclear lysates were clarified by centrifugation at 15100g for 10 minutes.

### Preparation of meiotic spreads from testicular cells

Meiotic spreads were prepared according to the published protocol (71).

### ChIP-sequencing of linker histone variant H1t in P20 mouse testicular cells

Chromatin immunoprecipitation (ChIP) was carried out using the published protocol (72). Briefly, P20 mice testes were dissected in 1X PBS (with 1% formaldehyde), incubated for 10 min on a rotating wheel at room temperature. Quenching to remove formaldehyde was done using glycine (250mM final concentration). The pellet was washed with 1X PBS multiple times. The suspension was filtered using a 40µm filter and then centrifuged at 500g for 5 min at 4°C. The pellet was resuspended in Buffer A (10mM Tris pH 8.0; 10mM KCL; 0.25% Triton-X-100; 1mM EDTA; 0.5mM EGTA; 1X protease inhibitor cocktail from Roche), and incubated on ice for 5 min. Again, the contents were centrifuged at 500g for 5 min at 4°C. The pellet was then resuspended in Buffer B (10mM Tris pH 8.0; 200mM NaCl; 1mM EDTA, 0.5mM EGTA; 1X protease inhibitor cocktail), incubated on ice for 10 min, centrifuged again at 500g for 5 min at 4°C. Finally, the pellet was resuspended in SDS lysis buffer (1% SDS, 10mM EDTA, 50mM Tris pH 8.0, 1X protease inhibitor cocktail). The contents were incubated on a rotating wheel for 30 min at 4°C. Sonication was carried out 40 cycles, and chromatin fractions were diluted 1/10 times with the dilution buffer (5mM Tris PH 8.0, 140mM NaCl, 0.5% TritonX-100, 0.05% sodium deoxycholate, 0.5mM EGTA) before subjecting it to antibody binding overnight at 4°C on a rotating wheel. Dynabeads were added the next day, washes were given with W1 (10mM Tris pH 8.0, 150mM KCl, 0.50% NP-40, 1mM EDTA), W2 (10mM Tris pH 8.0, 100mM NaCl, 0.10% sodium deoxycholate, 0.50% Triton-X-100), W3a (10mM Tris pH 8.0, 400mM NaCl, 0.10% sodium deoxycholate, 0.50% Triton-X-100), W3b (10mM Tris pH 8.0, 500mM NaCl, 0.10% sodium deoxycholate, 0.50% Triton-X-100), W4 (10mM Tris pH 8.0, 250mM LiCl, 0.50% sodium deoxycholate, 0.50% NP-40, 1mM EDTA), W5 (10mM Tris pH 8.0, 1mM EDTA) wash buffers. The complexes were eluted from input, rabbit IgG control ChIP, and H1t ChIP samples using elution buffer (50mM Tris pH 8.0, 1% SDS, 1mM EDTA) and then incubated at 65°C overnight with intermittent shaking. The supernatants recovered from the tubes were subjected to RNase A and proteinase K digestions at 37°C. DNA was then extracted from the elute fractions using the phenol-chloroform method. The input and ChIP DNA libraries were prepared using the NEBNext Ultra II DNA library preparation kit. The resulting libraries were quantified before getting sequenced on the Illumina HiSeqX system to generate 2X150 bp sequence reads.

### Computational data analysis

All the four samples (input controls two and H1t ChIP two samples - the raw Illumina sequenced files in FASTQ) were quality checked with FastQC (version 0.11.5). Each sample (two Input controls and two H1t ChIP files) were then aligned to mouse genome version (GRCm38.p5) using Bowtie2 (73) with alignment parameters: -D 15 -R 2 -N 1 -i S,1,0.75 -- local. The aligned samples (SAM/BAM files) were used to find the peaks with MACS2 version 2.1.2, with parameters: #effective genome size = 1.87e+09, #band width = 300 #model fold = (5, 50), #pvalue cutoff for narrow/strong regions = 5.00e-02, #pvalue cutoff for broad/weak regions = 5.00e-02, #Range for calculating regional lambda is: 1000 bp and 10000 bp, #Broad region calling is on, #Paired-End mode is on. The ChIP-sequencing dataset is given in Additional File 1. Annotation of H1t-bound genomic regions was carried out using HOMER (Additional File 2).

Various in house Perl scripts and database comparision scripts at BioCOS Life Sciences were used to do the rest of the analysis like (1) comparison with various datasets, rDNA, Motif, and comparative methylation analysis. The read concentration of H1t peaks was compared against the already published datasets of active gene promoters (GSE93955), TSS (GENCODE), DSB hotspots (GSE93955) and ATAC sequencing data (GSE102954) using in house scripts and software at BioCOS Life Sciences and plotted using ngsplot.r tool (74). FASTA sequences corresponding to the rDNA element were obtained from NCBI (40). A FASTQ file was generated from the significant peaks obtained from the MACS analysis to enable the alignment of the peaks on to rDNA FASTA sequences. After the peaks were aligned to the rDNA element using Bowtie using the rDNA FASTA as reference file, the tag densities were then computed on the various regions of the rDNA element (Fig 4A).

The raw bisulfite sequencing data for P20 mouse testicular cells were obtained from the Sequence Read Archive (SRA) database (SRS557654). SRR1170580 is the Single end WT sample from SRS557654, is used for extraction of methylation locations and compare locations with H1t Peaks. The reads were aligned using Bowtie2 onto the reference genome of mouse (GRCm38.p5)., The methylation analysis was done using bismark_v0.22.1 (75) and the methylated markers were isolated. (./bismark_v0.22.1/bismark --parallel 4 --bowtie2 -D 5 -R 1 -N 0 -L 22 --local -un -o AlignmentMeth_SE_SRS557654 --prefix AlignMethyl Bismark Genome mm10/single_end../Raw_SE_sample_SRS557654/SRR1170580.fastq.bz2). The methylated markers were isolated (./bismark_v0.22.1/bismark_methylation_extractor -s --comprehensive -o Methylated_Regions/ --cytosine_report --bedGraph --parallel 2 AlignmentMeth_SE_SRS557654/AlignMethylBismark.SRR1170580_bismark_bt2.bam -- genome_folder Genome_mm10/). Then the overlapping of the methylated markers (bases) of the database (SRS557654) with the H1t ChIP-seq Peaks was carried out using in house perl scripts and R (https://www.r-project.org/) (Fig 4B).

Motif analysis was carried out using MEME software (76). The parameters set for MEME-Search-size → 100000, DataBase: JASPAR/JASPAR2018_CORE_non-redundant.meme,- ccut(cut sequence from center): 0, -meme-searchsize: 100000,-filter-thresh: 0.0001(e Value),-meme-minw: 6,-meme-maxw: 50,-meme-p: 6,-meme, nmotifs: 15,-dreme-e: 0.0001,- dreme-m: 15,-centrimo-ethresh: 0.0001.

### Mass spectrometric identification of proteins associated with H1t-containing chromatin fragments

Immunoprecipitation of H1t-associated protein complexes was carried out, and the proteins were extracted from the beads using the elution buffer of the Pierce co-IP kit. The eluted proteins were resolved on 15% SDS gel and the gel was subjected to Coomassie staining. The stained wells corresponding to the IgG and IP lanes were outsourced for mass spectrometry to identify the interacting proteins. The detailed experimental protocol of the mass spectrometry was followed according to the published protocol (77). The list of H1t-associated proteins is given in Additional File 3. The comprehensive list of common proteins between H1t and the heterochromatin set (43) is given in Additional File 4.

### Western blotting

The proteins were resolved by gel electrophoresis and transferred onto the nitrocellulose membrane. The blots were blocked using blocking solution (5% skim milk in 1X TBS). The primary antibodies were made in 1% skim milk (prepared in 1X TBS) and added onto the blots and were left for overnight incubation in shaking conditions at 4°C. Next day, two washes were given using 0.1%TBST (0.1% Tween-20 in 1X TBS) for 5 minutes each. The secondary antibodies prepared in 1% skim milk was added to the blots and left for shaking for 1-hour incubation at room temperature. 2-3 washes were given with 0.1% TBST before developing the blot using Millipore Immobilon Forte Western HRP Substrate kit (Cat number-WBLUF0100) in the Biorad Chemidoc Touch imaging system.

For stripping of proteins from the blot, 1X Millipore Reblot Plus Strong Antibody Stripping Solution (Cat number-2504) was added to the blot and left for shaking at room temperature for 20-25 min. The blot was developed using the ECL kit to check for any signal on the membrane. Two washes were then given with blocking solution (5% skim milk in 1X TBS) for 5 min each and then proceeded with incubation with the primary antibody solution.

List of antibodies used in the present study-.

**List of antibodies used in the present study-**
| **Reagents** |  |  |  |  |
| --- | --- | --- | --- | --- |
| **Antibodies-** | **Host** | **Company Name** | **Cat number** | **Application** |
| Scp3 | Mouse | Abcam | ab97672 | I.F, W.B |
| HP1β | Mouse | Millipore | MAB3448 | W.B |
| HP1γ | Mouse | Millipore | 05-690 | W.B |
| Trim28 (Kap1) | Mouse | Abcam | ab22553 | W.B |
| Hormad1 | Rabbit | Abcam | ab155176 | W.B |
| NPM1 | Mouse | Abcam | ab10530 | W.B |
| Hspa2 | Rabbit | Invitrogen | PA5-81895 | W.B |
| PIWIL1 | Rabbit | Invitrogen | PA5-21051 | W.B |
| H3K9me3 | Rabbit | Millipore | 07-442 | W.B, ChIP |
| H4K20me3 | Rabbit | Abcam | ab9053 | W.B, ChIP |
| H1.2 | Rabbit | Abcam | ab181973 | W.B |

*W.B- Western Blotting; ChIP- Chromatin Immunoprecipitation; I.F- Immunofluorescence*

## List of Abbreviations

DDR: DNA double-strand break repair
TSS: Transcription Start Sites
LTR: Long Terminal Repeats
LINE: Long Interspersed Nuclear Element
SINE: Short Interspersed Nuclear Element
TE: Transposable Element
rDNA: Ribosomal DNA
ChIP: Chromatin Immunoprecipitation
ATAC-seq: Assay for Transposase-Accessible Chromatin using sequencing
piRNA: PIWI-interacting RNA
PIWI: P-element induced wimpy testis

## Declarations

### Ethics Approval

This work was approved by the Animal Ethics Committee of JNCASR, Bangalore, India.

### Consent for publication

Not applicable

### Availability of data and materials

The ChIP-sequencing dataset containing the raw and processed files are currently under submission to the Gene Expression Omnibus (GEO). The accession number will be provided once received.

### Author Contributions

I.A.M performed the experiments. I.A.M and M.R.S designed the experiments. I.A.M, S.K and M.R.S wrote the manuscript. S.K performed the computational data analysis associated with the H1t ChIP-sequencing dataset like overlap analysis with datasets, rDNA analysis, motif analysis, bisulphite sequencing etc. All authors discussed the results and approved the final manuscript.

## Acknowledgements

We acknowledge Dr. R.G Prakash of the Animal Facility and Suma B.S of the Confocal Imaging Facility at JNCASR. We also thank members of the Imgenex facility for carrying out the antibody generation project. We also thank Tomaino Ross of TAPLIN Mass spectrometry facility at Harvard for mass spectrometry based identification of interacting proteins.

## Table of Contents (Additional Figures)

**Additional Figure 1.**
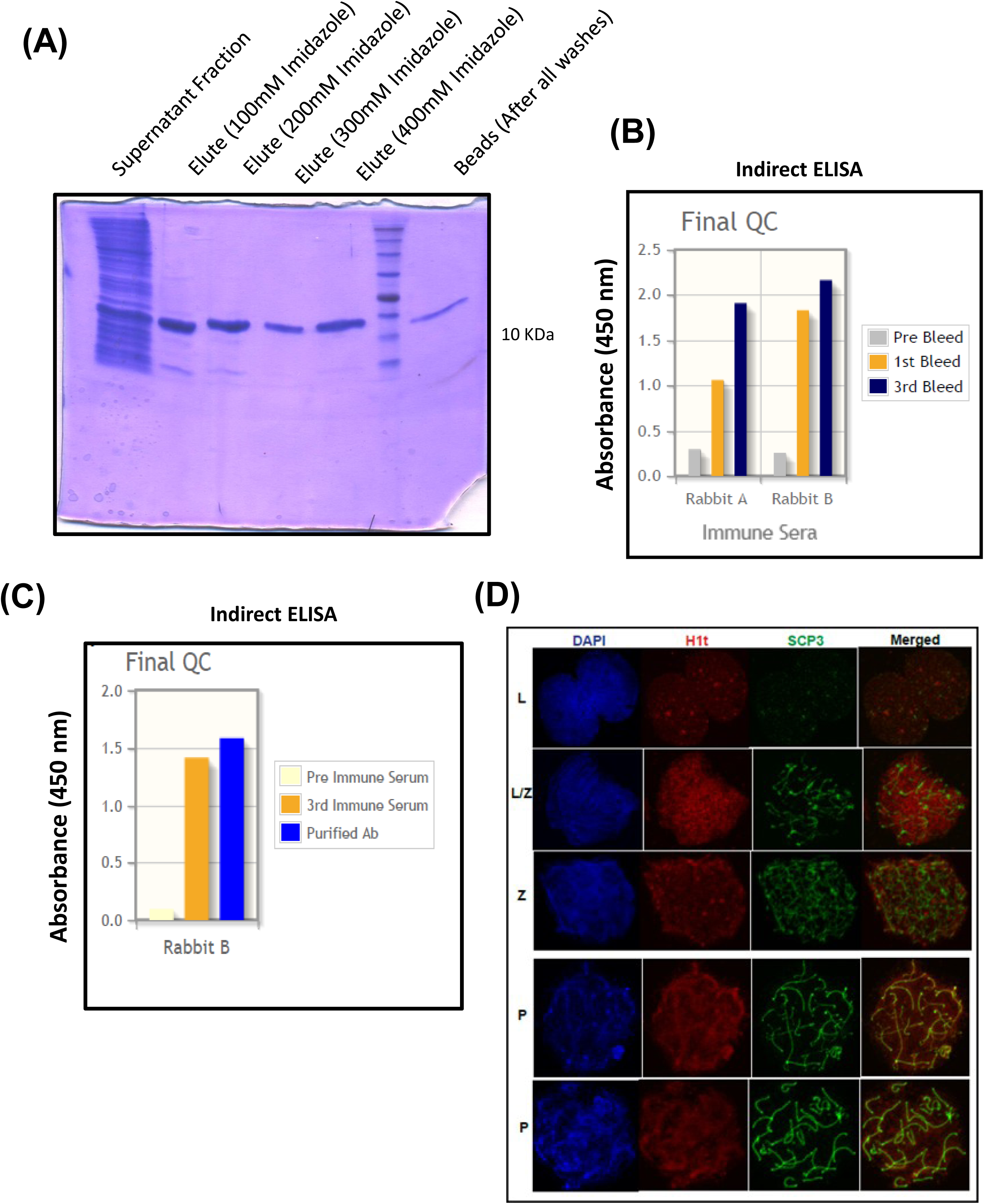
**A.** Coomassie-stained gel showing the successful purification of His tagged C-terminal fragment of H1t. The purity of proteins was determined after elution using 100mM, 200mM, 300mM, and 400mM imidazole, wherein the protein fractions were obtained after elution using 300/400mM concentration of imidazole. (**B**-**C)** Validation of specificity of the H1t antibody towards the recombinant H1t C-terminal protein fragment by ELISA using B. Immune sera and C. Purified antibody. The sera, as well as purified antibodies, showed reactivity against the H1t C-terminal protein fragment. The color code schemes have been indicated on the right of the figures. **D**. Immunostaining pattern of linker histone variant H1t across various stages of meiotic prophase I. Staining of anti-H1t and anti-Scp3 across leptotene (L, first panel), leptotene-zygotene (L/Z, second panel), zygotene (Z, third panel), and pachytene (P, fourth and fifth panels).

**Additional Figure 2.**
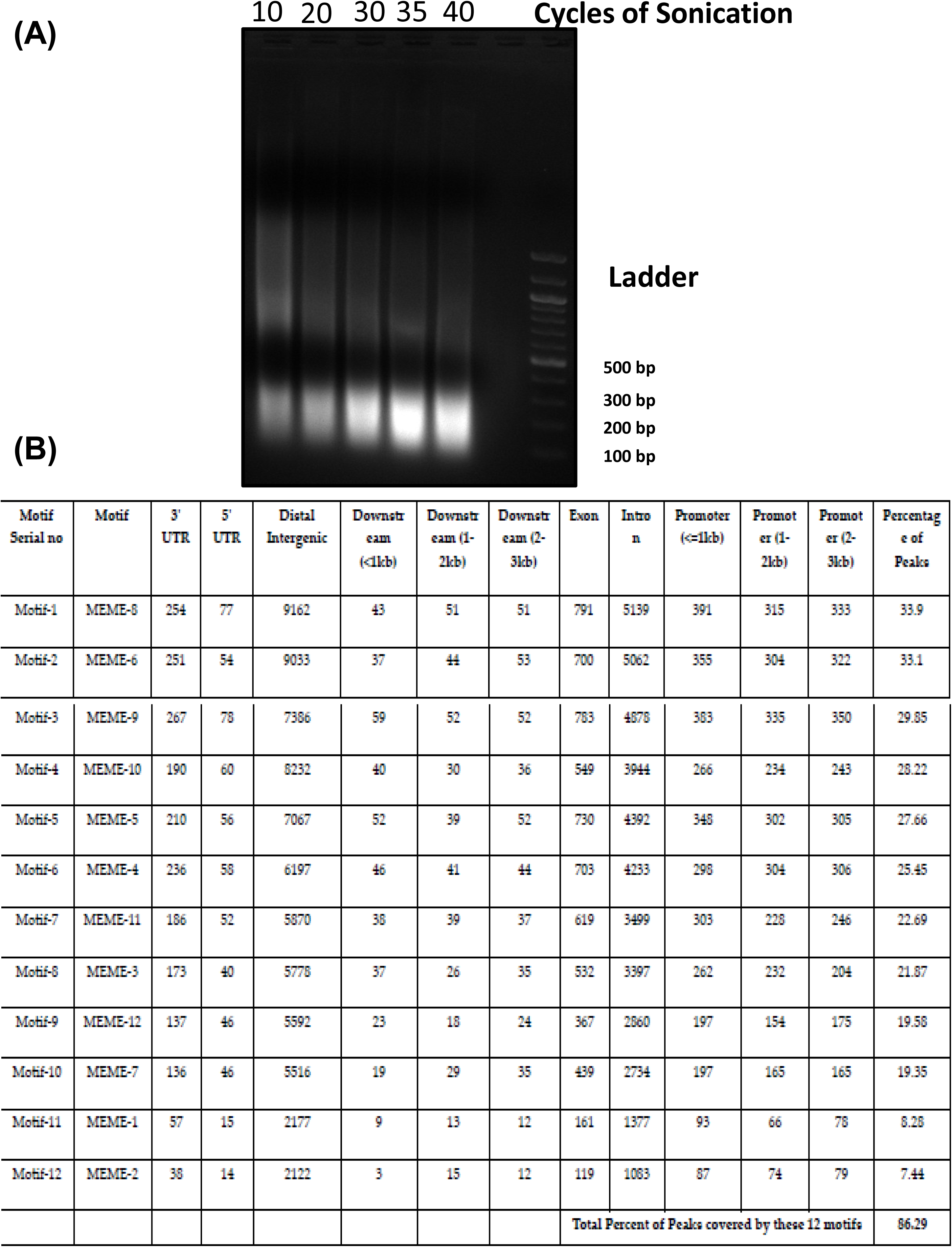
**A.** Profile of DNA fragments obtained after 10, 20, 30, 35, and 40 cycles of sonication of P20 mouse testicular chromatin. 100-300 bp of fragment sizes predominantly obtained after 40 cycles of sonication were used further for ChIP assays. **B.** Table of motifs identified of H1t bound genomic regions in pachytene spermatocytes using MEME software.

## Additional File Legends

**Additional File 1-** ChIP-sequencing peaks of H1t in P20 mouse testicular cells

**Additional File 2-** Annotation of H1t peaks using HOMER

**Additional File 3-** H1t-associated proteins obtained after mass spectrometry

**Additional File 4-** H1t and associated heterochromatin-related proteins

## References

1. Bharath MM, Chandra NR, Rao MR. Molecular modeling of the chromatosome particle. Nucleic Acids Res. 2003;31(14):4264–74.

2. Simpson RT. Structure of the chromatosome, a chromatin particle containing 160 base pairs of DNA and all the histones. Biochemistry. 1978;17(25):5524–31.

3. Bradbury EM, Chapman GE, Danby SE, Hartman PG, Riches PL. Studies on the role and mode of operation of the very-lysine-rich histone H1 (F1) in eukaryote chromatin. The properties of the N-terminal and C-terminal halves of histone H1. Eur J Biochem. 1975;57(2):521–8.

4. Hartman PG, Chapman GE, Moss T, Bradbury EM. Studies on the role and mode of operation of the very-lysine-rich histone H1 in eukaryote chromatin. The three structural regions of the histone H1 molecule. Eur J Biochem. 1977;77(1):45–51.

5. Aviles FJ, Danby SE, Chapman GE, Crane-Robinson C, Bradbury EM. The conformation of histone H5 bound to DNA. Maintenance of the globular structure after binding. Biochim Biophys Acta. 1979;578(2):290–6.

6. Rattle HW, Langan TA, Danby SE, Bradbury EM. Studies on the role and mode of operation of the very-lysine-rich histones in eukaryote chromatin. Effect of A and B site phosphorylation on the conformation and interaction of histone H1. Eur J Biochem. 1977;81(3):499–505.

7. Roque A, Ponte I, Suau P. Interplay between histone H1 structure and function. Biochim Biophys Acta. 2016;1859(3):444–54.

8. Hendzel MJ, Lever MA, Crawford E, Th’ng JP. The C-terminal domain is the primary determinant of histone H1 binding to chromatin in vivo. J Biol Chem. 2004;279(19):20028–34.

9. Lennox RW, Cohen LH. The alterations in H1 histone complement during mouse spermatogenesis and their significance for H1 subtype function. Dev Biol. 1984;103(1):80–4.

10. Steger K, Klonisch T, Gavenis K, Drabent B, Doenecke D, Bergmann M. Expression of mRNA and protein of nucleoproteins during human spermiogenesis. Mol Hum Reprod. 1998;4(10):939–45.

11. Drabent B, Bode C, Bramlage B, Doenecke D. Expression of the mouse testicular histone gene H1t during spermatogenesis. Histochem Cell Biol. 1996;106(2):247–51.

12. Drabent B, Bode C, Miosge N, Herken R, Doenecke D. Expression of the mouse histone gene H1t begins at premeiotic stages of spermatogenesis. Cell Tissue Res. 1998;291(1):127–32.

13. Bucci LR, Brock WA, Meistrich ML. Distribution and synthesis of histone 1 subfractions during spermatogenesis in the rat. Exp Cell Res. 1982;140(1):111–8.

14. Grimes SR, Wilkerson DC, Noss KR, Wolfe SA. Transcriptional control of the testis-specific histone H1t gene. Gene. 2003;304:13–21.

15. Govin J, Caron C, Lestrat C, Rousseaux S, Khochbin S. The role of histones in chromatin remodelling during mammalian spermiogenesis. Eur J Biochem. 2004;271(17):3459–69.

16. Lin Q, Sirotkin A, Skoultchi AI. Normal spermatogenesis in mice lacking the testis-specific linker histone H1t. Mol Cell Biol. 2000;20(6):2122–8.

17. Drabent B, Benavente R, Hoyer-Fender S. Histone H1t is not replaced by H1.1 or H1.2 in pachytene spermatocytes or spermatids of H1t-deficient mice. Cytogenet Genome Res. 2003;103(3-4):307–13.

18. Fantz DA, Hatfield WR, Horvath G, Kistler MK, Kistler WS. Mice with a targeted disruption of the H1t gene are fertile and undergo normal changes in structural chromosomal proteins during spermiogenesis. Biol Reprod. 2001;64(2):425–31.

19. Khadake JR, Rao MR. DNA- and chromatin-condensing properties of rat testes H1a and H1t compared to those of rat liver H1bdec; H1t is a poor condenser of chromatin. Biochemistry. 1995;34(48):15792–801.

20. De Lucia F, Faraone-Mennella MR, D’Erme M, Quesada P, Caiafa P, Farina B. Histone-induced condensation of rat testis chromatin: testis-specific H1t versus somatic H1 variants. Biochem Biophys Res Commun. 1994;198(1):32–9.

21. Suzuki M. SPKK, a new nucleic acid-binding unit of protein found in histone. EMBO J. 1989;8(3):797–804.

22. Drabent B, Kardalinou E, Doenecke D. Structure and expression of the human gene encoding testicular H1 histone (H1t). Gene. 1991;103(2):263–8.

23. Bharath MM, Chandra NR, Rao MR. Prediction of an HMG-box fold in the C-terminal domain of histone H1: insights into its role in DNA condensation. Proteins. 2002;49(1):71–81.

24. Khadake JR, Rao MR. Condensation of DNA and chromatin by an SPKK-containing octapeptide repeat motif present in the C-terminus of histone H1. Biochemistry. 1997;36(5):1041–51.

25. Ramesh S, Bharath MM, Chandra NR, Rao MR. A K52Q substitution in the globular domain of histone H1t modulates its nucleosome binding properties. FEBS Lett. 2006;580(25):5999–6006.

26. Nagamori I, Kobayashi H, Shiromoto Y, Nishimura T, Kuramochi-Miyagawa S, Kono T, et al. Comprehensive DNA Methylation Analysis of Retrotransposons in Male Germ Cells. Cell Rep. 2015;12(10):1541–7.

27. Shoji M, Tanaka T, Hosokawa M, Reuter M, Stark A, Kato Y, et al. The TDRD9-MIWI2 complex is essential for piRNA-mediated retrotransposon silencing in the mouse male germline. Dev Cell. 2009;17(6):775–87.

28. Ma L, Buchold GM, Greenbaum MP, Roy A, Burns KH, Zhu H, et al. GASZ is essential for male meiosis and suppression of retrotransposon expression in the male germline. PLoS Genet. 2009;5(9):e1000635.

29. Kojima K, Kuramochi-Miyagawa S, Chuma S, Tanaka T, Nakatsuji N, Kimura T, et al. Associations between PIWI proteins and TDRD1/MTR-1 are critical for integrated subcellular localization in murine male germ cells. Genes Cells. 2009;14(10):1155–65.

30. Reuter M, Chuma S, Tanaka T, Franz T, Stark A, Pillai RS. Loss of the Mili-interacting Tudor domain-containing protein-1 activates transposons and alters the Mili-associated small RNA profile. Nat Struct Mol Biol. 2009;16(6):639–46.

31. Tani R, Hayakawa K, Tanaka S, Shiota K. Linker histone variant H1T targets rDNA repeats. Epigenetics. 2016;11(4):288–302.

32. Maezawa S, Yukawa M, Alavattam KG, Barski A, Namekawa SH. Dynamic reorganization of open chromatin underlies diverse transcriptomes during spermatogenesis. Nucleic Acids Res. 2018;46(2):593–608.

33. Bourc’his D, Bestor TH. Meiotic catastrophe and retrotransposon reactivation in male germ cells lacking Dnmt3L. Nature. 2004;431(7004):96–9.

34. Molaro A, Falciatori I, Hodges E, Aravin AA, Marran K, Rafii S, et al. Two waves of de novo methylation during mouse germ cell development. Genes Dev. 2014;28(14):1544–9.

35. Pezic D, Manakov SA, Sachidanandam R, Aravin AA. piRNA pathway targets active LINE1 elements to establish the repressive H3K9me3 mark in germ cells. Genes Dev. 2014;28(13):1410–28.

36. Delaval K, Govin J, Cerqueira F, Rousseaux S, Khochbin S, Feil R. Differential histone modifications mark mouse imprinting control regions during spermatogenesis. EMBO J. 2007;26(3):720–9.

37. Grewal SI, Jia S. Heterochromatin revisited. Nat Rev Genet. 2007;8(1):35–46.

38. Peters AH, O’Carroll D, Scherthan H, Mechtler K, Sauer S, Schofer C, et al. Loss of the Suv39h histone methyltransferases impairs mammalian heterochromatin and genome stability. Cell. 2001;107(3):323–37.

39. Peters AH, Schubeler D. Methylation of histones: playing memory with DNA. Curr Opin Cell Biol. 2005;17(2):230–8.

40. Grozdanov P, Georgiev O, Karagyozov L. Complete sequence of the 45-kb mouse ribosomal DNA repeat: analysis of the intergenic spacer. Genomics. 2003;82(6):637–43.

41. Reuter M, Berninger P, Chuma S, Shah H, Hosokawa M, Funaya C, et al. Miwi catalysis is required for piRNA amplification-independent LINE1 transposon silencing. Nature. 2011;480(7376):264–7.

42. Saxe JP, Chen M, Zhao H, Lin H. Tdrkh is essential for spermatogenesis and participates in primary piRNA biogenesis in the germline. EMBO J. 2013;32(13):1869–85.

43. Guthmann M, Burton A, Torres-Padilla ME. Expression and phase separation potential of heterochromatin proteins during early mouse development. EMBO Rep. 2019:e47952.

44. Dong J, Wang X, Cao C, Wen Y, Sakashita A, Chen S, et al. UHRF1 suppresses retrotransposons and cooperates with PRMT5 and PIWI proteins in male germ cells. Nat Commun. 2019;10(1):4705.

45. Percharde M, Lin CJ, Yin Y, Guan J, Peixoto GA, Bulut-Karslioglu A, et al. A LINE1-Nucleolin Partnership Regulates Early Development and ESC Identity. Cell. 2018;174(2):391–405 e19.

46. J. Yuyang Lu LC, Tong Li, Ting Wang, Yafei Yin, Ge Zhan, Ke Zhang, Michelle Percharde, Liang Wang, Qi Peng, Pixi Yan, Hui Zhang, Xue Han, Xianju Bi, Wen Shao, Yantao Hong, Zhongyang Wu, Peizhe Wang, Wenzhi Li, Jing Zhang, Zai Chang, Yingping Hou, Pilong Li, Miguel Ramalho-Santos, Jie Na, Wei Xie, Yujie Sun, Xiaohua Shen. L1 and B1 repeats blueprint the spatial organization of chromatin. BioRxiv. 2019.

47. Zamudio N, Barau J, Teissandier A, Walter M, Borsos M, Servant N, et al. DNA methylation restrains transposons from adopting a chromatin signature permissive for meiotic recombination. Genes Dev. 2015;29(12):1256–70.

48. Fan Y, Nikitina T, Zhao J, Fleury TJ, Bhattacharyya R, Bouhassira EE, et al. Histone H1 depletion in mammals alters global chromatin structure but causes specific changes in gene regulation. Cell. 2005;123(7):1199–212.

49. Choi J, Lyons DB, Kim MY, Moore JD, Zilberman D. DNA Methylation and Histone H1 Jointly Repress Transposable Elements and Aberrant Intragenic Transcripts. Mol Cell. 2019.

50. Iwasaki YW, Murano K, Ishizu H, Shibuya A, Iyoda Y, Siomi MC, et al. Piwi Modulates Chromatin Accessibility by Regulating Multiple Factors Including Histone H1 to Repress Transposons. Mol Cell. 2016;63(3):408–19.

51. Zentner GE, Balow SA, Scacheri PC. Genomic characterization of the mouse ribosomal DNA locus. G3 (Bethesda). 2014;4(2):243–54.

52. Ghosh AK, Hoff CM, Jacob ST. Characterization of the 130-bp repeat enhancer element of the rat ribosomal gene: functional interaction with transcription factor E1BF. Gene. 1993;125(2):217–22.

53. Cong R, Das S, Douet J, Wong J, Buschbeck M, Mongelard F, et al. macroH2A1 histone variant represses rDNA transcription. Nucleic Acids Res. 2014;42(1):181–92.

54. Bickmore WA, van Steensel B. Genome architecture: domain organization of interphase chromosomes. Cell. 2013;152(6):1270–84.

55. Bickmore WA. The spatial organization of the human genome. Annu Rev Genomics Hum Genet. 2013;14:67–84.

56. Dekker J, Marti-Renom MA, Mirny LA. Exploring the three-dimensional organization of genomes: interpreting chromatin interaction data. Nat Rev Genet. 2013;14(6):390–403.

57. Belmont AS. Large-scale chromatin organization: the good, the surprising, and the still perplexing. Curr Opin Cell Biol. 2014;26:69–78.

58. Pombo A, Dillon N. Three-dimensional genome architecture: players and mechanisms. Nat Rev Mol Cell Biol. 2015;16(4):245–57.

59. Sexton T, Cavalli G. The role of chromosome domains in shaping the functional genome. Cell. 2015;160(6):1049–59.

60. Nemeth A, Conesa A, Santoyo-Lopez J, Medina I, Montaner D, Peterfia B, et al. Initial genomics of the human nucleolus. PLoS Genet. 2010;6(3):e1000889.

61. van Koningsbruggen S, Gierlinski M, Schofield P, Martin D, Barton GJ, Ariyurek Y, et al. High-resolution whole-genome sequencing reveals that specific chromatin domains from most human chromosomes associate with nucleoli. Mol Biol Cell. 2010;21(21):3735–48.

62. Dillinger S, Straub T, Nemeth A. Nucleolus association of chromosomal domains is largely maintained in cellular senescence despite massive nuclear reorganisation. PLoS One. 2017;12(6):e0178821.

63. Carone BR, Hung JH, Hainer SJ, Chou MT, Carone DM, Weng Z, et al. High-resolution mapping of chromatin packaging in mouse embryonic stem cells and sperm. Dev Cell. 2014;30(1):11–22.

64. Samans B, Yang Y, Krebs S, Sarode GV, Blum H, Reichenbach M, et al. Uniformity of nucleosome preservation pattern in Mammalian sperm and its connection to repetitive DNA elements. Dev Cell. 2014;30(1):23–35.

65. Mishra LN, Shalini V, Gupta N, Ghosh K, Suthar N, Bhaduri U, et al. Spermatid-specific linker histone HILS1 is a poor condenser of DNA and chromatin and preferentially associates with LINE-1 elements. Epigenetics & Chromatin. 2018;11.

66. Faraone-Mennella MR, De Lucia F, Gentile N, Quesada P, Farina B. In vitro poly(ADP-ribosyl)ated histones H1a and H1t modulate rat testis chromatin condensation differently. J Cell Biochem. 1999;76(1):20–9.

67. Meyer-Ficca ML, Ihara M, Lonchar JD, Meistrich ML, Austin CA, Min W, et al. Poly(ADP-ribose) Metabolism Is Essential for Proper Nucleoprotein Exchange During Mouse Spermiogenesis. Biology of Reproduction. 2011;84(2):218–28.

68. Luense LJ, Wang X, Schon SB, Weller AH, Lin Shiao E, Bryant JM, et al. Comprehensive analysis of histone post-translational modifications in mouse and human male germ cells. Epigenetics Chromatin. 2016;9:24.

69. Montellier E, Boussouar F, Rousseaux S, Zhang K, Buchou T, Fenaille F, et al. Chromatin-to-nucleoprotamine transition is controlled by the histone H2B variant TH2B. Genes Dev. 2013;27(15):1680–92.

70. Somyajit K, Basavaraju S, Scully R, Nagaraju G. ATM- and ATR-mediated phosphorylation of XRCC3 regulates DNA double-strand break-induced checkpoint activation and repair. Mol Cell Biol. 2013;33(9):1830–44.

71. Peters AH, Plug AW, van Vugt MJ, de Boer P. A drying-down technique for the spreading of mammalian meiocytes from the male and female germline. Chromosome Res. 1997;5(1):66–8.

72. Tardat M, Brustel J, Kirsh O, Lefevbre C, Callanan M, Sardet C, et al. The histone H4 Lys 20 methyltransferase PR-Set7 regulates replication origins in mammalian cells. Nat Cell Biol. 2010;12(11):1086–93.

73. Langmead B, Salzberg SL. Fast gapped-read alignment with Bowtie 2. Nat Methods. 2012;9(4):357–9.

74. Shen L, Shao N, Liu X, Nestler E. ngs.plot: Quick mining and visualization of next-generation sequencing data by integrating genomic databases. BMC Genomics. 2014;15:284.

75. Krueger F, Andrews SR. Bismark: a flexible aligner and methylation caller for Bisulfite-Seq applications. Bioinformatics. 2011;27(11):1571–2.

76. Bailey TL, Williams N, Misleh C, Li WW. MEME: discovering and analyzing DNA and protein sequence motifs. Nucleic Acids Res. 2006;34(Web Server issue):W369–73.

77. Mahadevan IA, Pentakota S, Roy R, Bhaduri U, Satyanarayana Rao MR. TH2BS11ph histone mark is enriched in the unsynapsed axes of the XY body and predominantly associates with H3K4me3-containing genomic regions in mammalian spermatocytes. Epigenetics Chromatin. 2019;12(1):53.

